# Immune Organization is Encoded in Coordinated Transcriptional Variation Across Cell Types

**DOI:** 10.64898/2026.03.10.710922

**Authors:** H. Jabran Zahid

## Abstract

Immune responses are often interpreted as cell type-specific transcriptional programs driven by external stimuli, with variation typically analyzed within each cell type. Here we show that immune organization is reflected in coordinated changes in gene expression across cell types within individuals, forming a shared low-dimensional state space. Using single-cell transcriptomes from a large longitudinal cohort, we find that thousands of genes vary coherently relative to subtype-specific reference profiles, defining this low-dimensional structure. Across individuals and over time, these deviations organize into a small number of recurring, temporally stratified patterns of transcriptional change. Dominant patterns reflect temporal dynamics shared across individuals, whereas others encode stable individual-specific transcriptional configurations. This donor-level coordination is observed across tissues, is reflected in cell surface protein expression and is captured in matched serum proteomics. Importantly, responses to in vitro cytokine stimulation are coordinated across cell types within donors, demonstrating that donor-level transcriptional organization is preserved under perturbation. Together, these results show that donor-level organization is evident in both transcriptional state and responses to perturbation, manifesting across cell types, tissues and time.

## Introduction

The immune system coordinates diverse cell types and compartments to sustain host defense while maintaining tolerance and homeostasis [1, 2, 3, 4]. Perturbations engage multiple immune compartments, indicating coordinated responses across the system [5, 6, 7]. This organization is often characterized at a high level through cell-type composition, intercellular signaling networks and circulating measurements [8, 4]. Single-cell transcriptional analyses, however, typically organize variation around cell identities, cell states and cell-type-specific programs [9, 10]. Prior work has established that individuals exhibit coordinated immune phenotypes and baseline immune states across multiple modalities. However, it remains unclear whether this donor-level organization is reflected in transcription. Here we show that this organization is directly observable in coordinated transcriptional variation across cell types.

Recent advances in high-dimensional immune profiling, including single-cell transcriptomics, proteomics and mass cytometry, enable systematic characterization of immune cell populations at scale [11, 8, 12]. In particular, single-cell RNA sequencing provides a direct readout of gene expression across thousands of genes in individual cells [13, 14, 9]. While these approaches have revealed extensive cellular heterogeneity [10], single-cell analyses typically treat inter-individual variation as context or a covariate rather than as a primary source of structure. As a result, coordinated system-level structure in transcription has not been directly characterized despite the availability of high-dimensional measurements.

Detecting shared system-level structure across heterogeneous immune cell populations requires repeated sampling within individuals and integration across cell types, yet most single-cell studies are cross-sectional and treat inter-individual variation as a secondary effect. As a result, whether transcription directly encodes system-level immune organization and whether that organization is preserved under perturbation, has remained unclear. Recent large-scale longitudinal and perturbation datasets [15, 16, 17, 18, 19] now enable direct testing of these questions.

We test whether system-level immune organization is reflected in transcription using single-cell transcriptomes from a large longitudinal cohort of healthy adults sampled repeatedly over time [19]. This cohort was previously used to characterize age-related immune dynamics and vaccine responses through integrated multiomic profiling. Here we quantify gene expression deviations relative to cell type-specific reference profiles and show that these deviations are coordinated across immune cell types within individuals. This shared structure spans major immune cell types and defines a low-dimensional state space captured by a small number of patterns of transcriptional change across donors and over time. This organization is consistent with the coexistence of stable inter-individual differences and within-individual dynamics observed in large cohort studies [20, 21, 22, 23]. It extends across tissues and is reflected in cell surface protein expression and in matched serum proteomics; importantly, it is also preserved under perturbation.

## Results

The Sound Life cohort comprises peripheral blood samples from 96 healthy adults (49 young, 25-35 years; 47 older, 55-65 years) collected at multiple visits across two annual cycles, enabling longitudinal analyses of immune transcription within donors. Immune cells are annotated at high resolution into 71 immune cell subsets, including major lineages such as T cells, B cells, NK cells and monocytes [19]. This cohort structure enables direct examination of coordinated transcriptional variation across immune cell types at the donor level.

To examine donor-level immune organization, our analysis proceeds in three biologically motivated steps (Fig. 1). First, we summarize gene expression within each immune subtype for every donor and visit by averaging expression profiles across all cells assigned to that subtype. This defines mean donor-visit-subtype transcriptional profiles for each donor and visit. Second, we remove subtype-specific reference profiles by centering profiles within each subtype relative to the cohort mean expression profile for that subtype. This step isolates donor-level transcriptional variation by removing the component associated with subtype identity. Third, we examine whether these deviations are coordinated across immune cell types and whether they define a shared transcriptional space that captures donor-level immune organization. Figure 1 schematically summarizes this analytical framework and the resulting structure of transcriptional variation.

**Figure 1:**
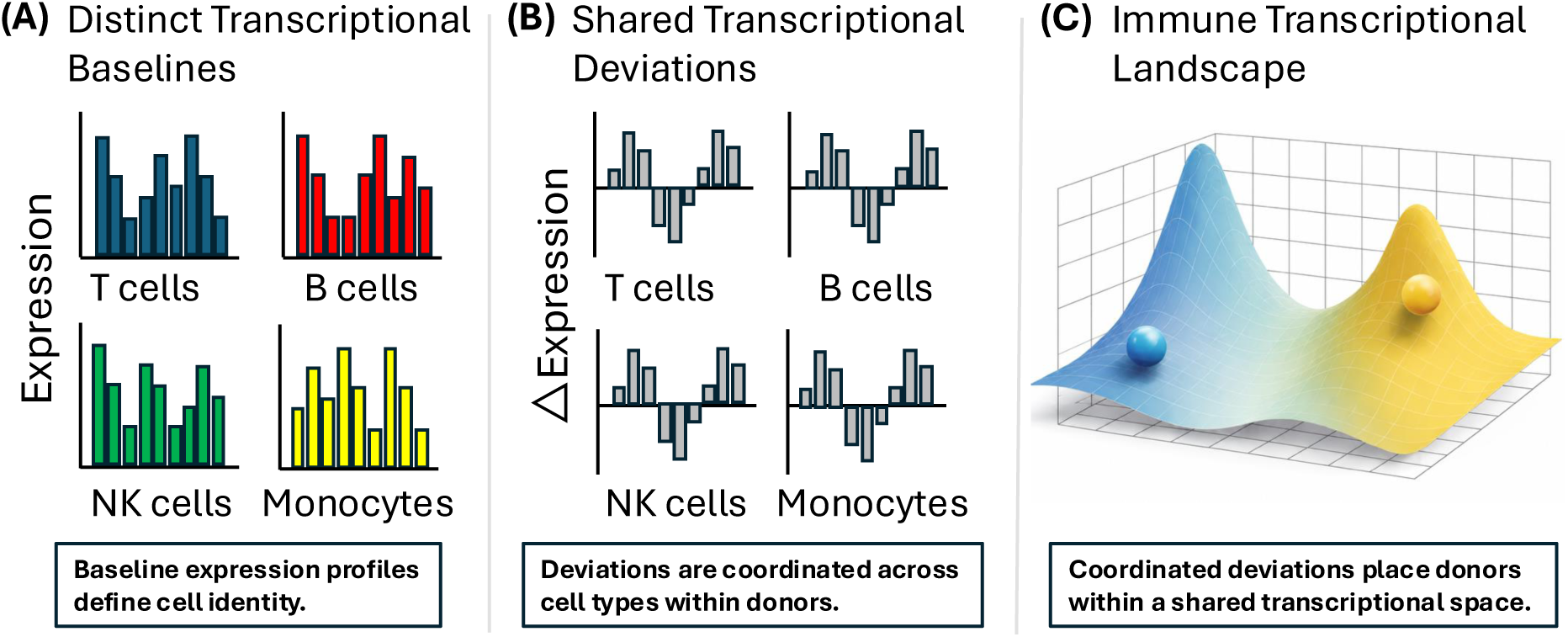
Conceptual and analytical overview of coordinated transcriptional variation across immune cell types. (A) Distinct gene expression profiles define immune cell identity across major immune lineages, illustrated as differing baseline transcriptional patterns for T cells, B cells, NK cells and monocytes. Gene expression is summarized for each donor, visit and subtype by averaging across cells to define donor-visit-subtype profiles. (B) To quantify how subtype-specific expression profiles vary across donors and over time, we measure deviations relative to a reference profile defined as the mean donor-visit-subtype profile for each subtype across all donor-visit instances in the cohort. This reference profile is subtracted gene-wise from each donor-visit-subtype profile, yielding deviation profiles that capture donor-specific variation at each visit. Within individuals, these deviations are strongly correlated across immune cell types, producing shared patterns of transcriptional change. (C) Deviation profiles are pooled across donors, visits and subtypes to define a low-dimensional transcriptional state space shared across individuals, which we refer to as the immune transcriptional landscape (ITL). Coordinated deviations place donors within this shared transcriptional space, providing a representation of coordinated transcriptional variation across immune cell types.

For each donor, visit and immune subtype, we compute a donor-visit-subtype profile by averaging log1p-normalized expression vectors across all cells assigned to that subtype. Groups containing fewer than 100 cells are excluded to ensure stable estimates. Analyses are restricted to a shared set of 3,530 genes defined by prevalence-based filtering across immune subtypes after exclusion of mitochondrial and ribosomal genes, enabling gene-wise comparison of transcriptional patterns across cell types. A small fraction of cells with strong transient activation, interferon-associated or cell-cycle signatures were removed before aggregation (see Methods). For each immune subtype, we define a reference transcriptional profile as the mean of donor–visit–subtype profiles across all donors and visits. Subtracting this reference from each donor–visit–subtype profile yields a deviation profile that captures gene expression variation relative to the characteristic transcriptional state of that subtype. This centering step isolates donor-level variation and is required for direct comparison across immune cell types. These deviation profiles serve as the fundamental units of all donor-level analyses.

### Coordinated transcriptional deviations across immune cell types

Immune cell types exhibit distinct transcriptional programs that dominate baseline gene expression differences across subtypes (Fig. 2A). After removing these cell type-specific reference profiles, we examine whether deviations are coordinated across immune cell types within individual donors. For an illustrative pair of immune subtypes, deviation profiles from the same donor are positively correlated across genes (Fig. 2B), indicating coordinated up- or down-regulation across cell types despite their distinct baseline transcriptional states. This alignment reflects donor-level coordination of transcriptional shifts across immune cell types; when sub-types are randomly paired across donors at the same visit the correlation is lost (Fig. 2C). Extending this analysis across T cells, B cells, NK cells and monocytes reveals consistently positive gene-wise correlations (Fig. 2D), demonstrating that coordinated transcriptional variation generalizes across immune cell types and donors. Although Fig. 2 illustrates a single donor–visit example, deviation profiles are positively correlated across subtype pairs in each donor–visit instance in the cohort, indicating that transcriptional variation is coordinated at the donor level.

**Figure 2:**
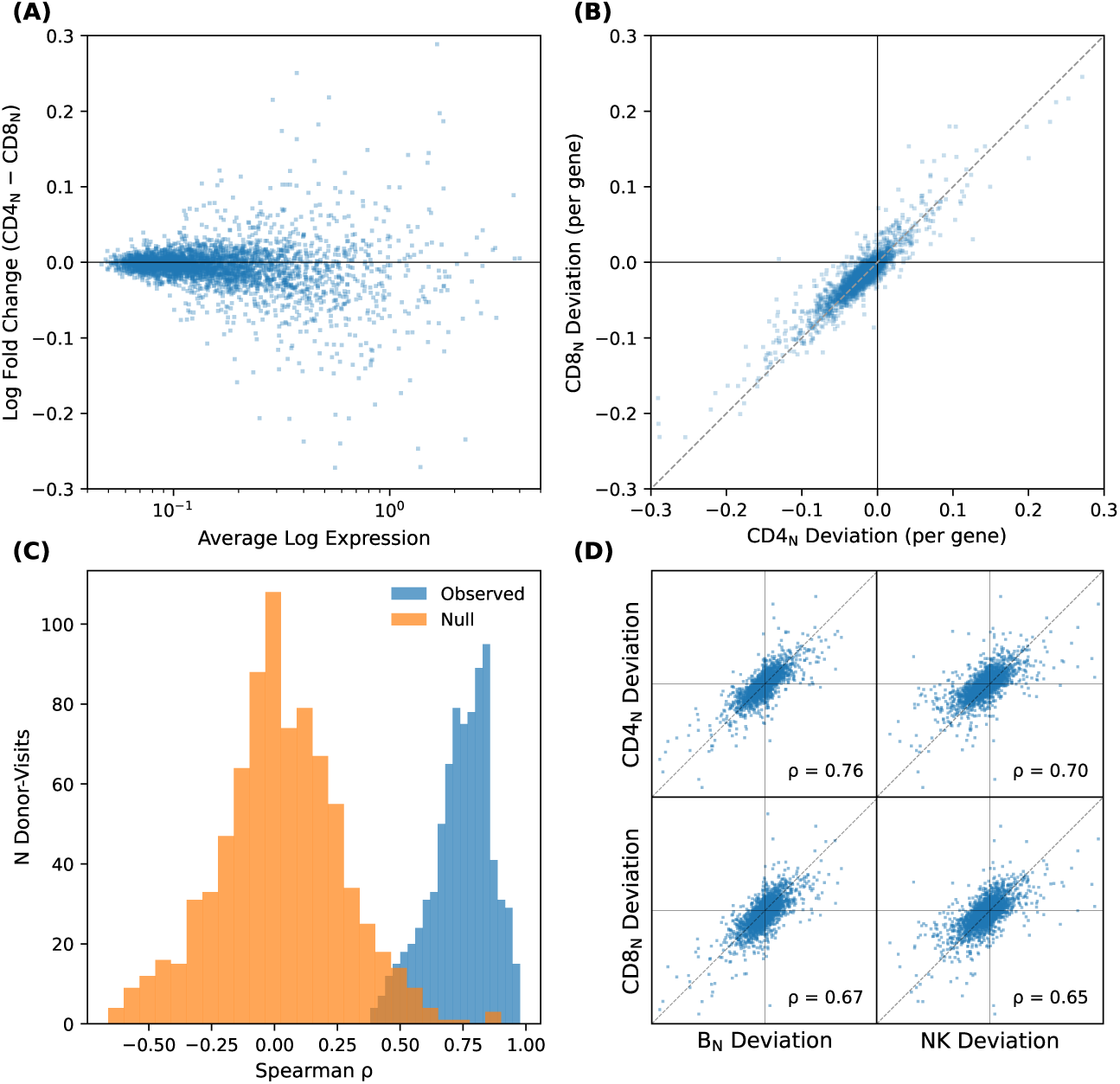
Coordinated transcriptional deviations across immune cell types. (A) Mean gene expression profiles differ systematically across cell types, reflecting cell type-specific baseline transcriptional programs. (B) After subtracting the mean gene expression profile for each cell type, deviations in expression for each gene are positively correlated across cell types within the same donor. Each point corresponds to a single gene, with values representing expression relative to the cohort-average reference profile for that cell type. This indicates coordinated up- or down-regulation of genes across cell types. (C) Distribution of correlation coefficients across all donor-visit pairs compared with a null generated by shuffling donor identities within each visit (preserving visit context and forbidding self-matches), demonstrating that the observed alignment depends on within-donor correspondence. (D) Extending this analysis across additional immune cell types reveals consistent positive correlations between T cells, B cells, NK cells and monocytes. Together, these analyses show that after accounting for cell type-specific reference profiles, transcriptional deviations are systematically aligned across immune cell types within donors, indicating structured and coordinated variation rather than independent fluctuations. Comparisons in (A, B, D) are shown for an illustrative donor at a single visit; CD4 and CD8 naive cells are used for illustration in (A-C) and similar relationships are observed across all well-sampled subtypes.

To quantify how much of this coordinated variation reflects stable inter-donor differences versus temporal change within donors, we partition total subtype-resolved variance in deviation profiles into between-donor and within-donor components. Across subtypes, within-donor temporal variation accounts for approximately 55-65% of total variance. Conversely, stable between-donor differences account for approximately 35-45%.

These results demonstrate coordinated transcriptional variation across immune cell types within donors and show that temporally varying within-donor changes modestly exceed stable between-donor differences in magnitude. We next ask whether this variation reflects a shared donor-level structure.

### Shared donor-level axes of transcriptional variation

Donor-level transcriptional variation is coordinated across immune cell types. We assess whether this shared variation can be consolidated into a unified donor representation. To do so, we (i) assess consistency of signals across subtypes within each compartment and (ii) test concordance of donor-level signals across compartments. We then evaluate the temporal coherence of the resulting representation as a measure of its biological validity.

To assess consistency of donor-level signals across subtypes, we perform principal component analysis (PCA) independently within each compartment (see Methods). We project subtype-specific expression profiles onto these axes to obtain per-subtype scores for each donor–visit. These scores are highly concordant across subtypes within each compartment, indicating that they reflect repeated measurements of a common compartment-level signal despite differences in subtype identity. We therefore aggregate these scores within each donor and visit to obtain a single compartment-level donor representation.

To assess concordance of donor-level signals across compartments, we compare the compartment-level donor representations derived above. We first examine donor ordering along the first principal component computed independently within each compartment. Donor ordering along this component is strongly correlated across T cells, B cells, NK cells and monocytes at a fixed visit (Fig. 3A). This concordance indicates that the first principal component reflects a transcriptional axis shared across immune compartments. Projecting deviation profiles into a reference PCA basis (CD4 naive), we further show that this shared structure extends beyond the first principal component (Fig. 3B), supporting a multi-dimensional organization of transcriptional variation shared across compartments. These results demonstrate that deviations from subtype-specific reference profiles align across immune compartments at the donor level, providing a basis for averaging compartment-level principal component scores to obtain a consolidated donor representation.

**Figure 3:**
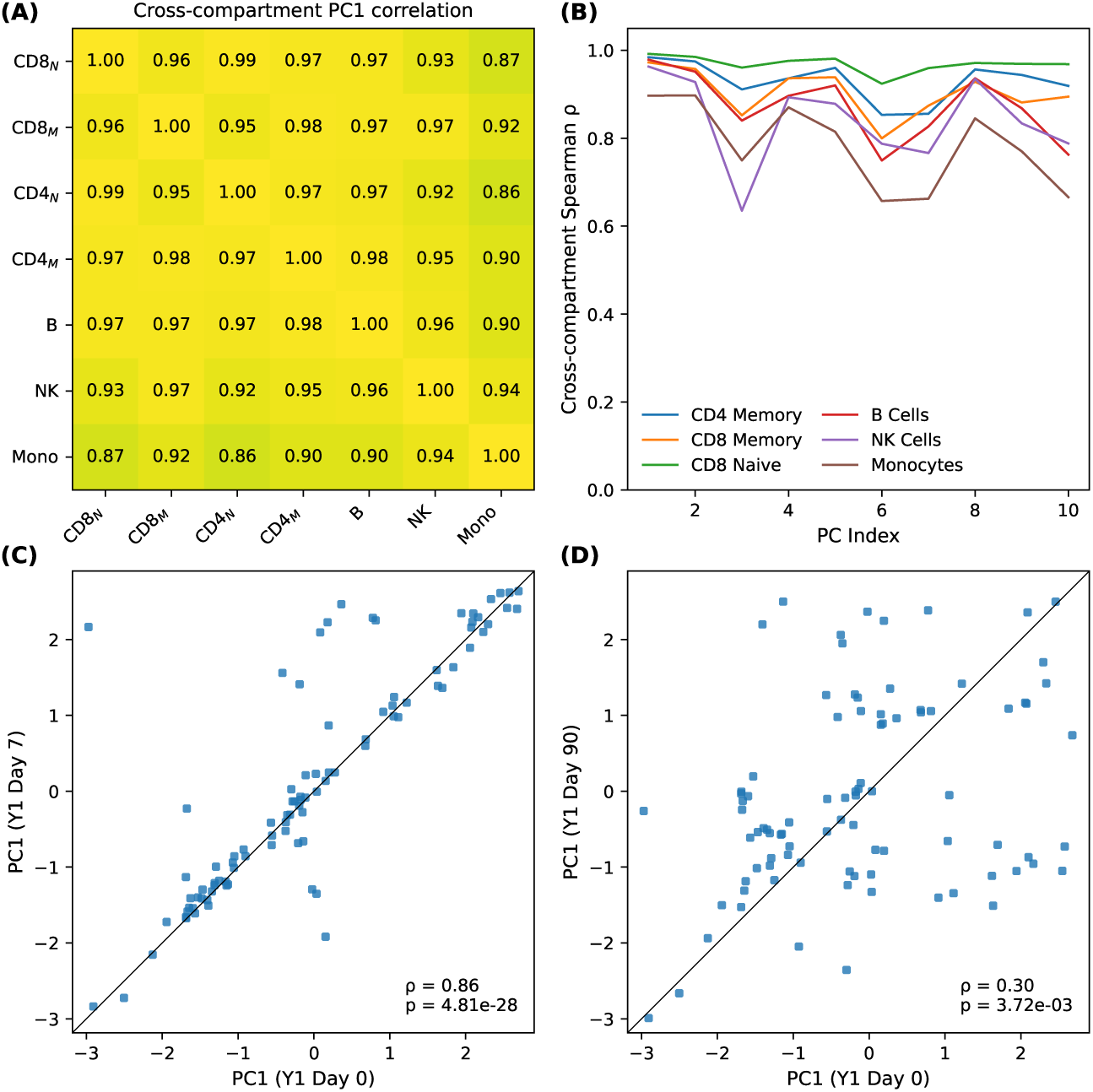
Independent recovery of shared donor-level transcriptional structure and its temporal coherence across immune compartments. (A) Principal component analysis (PCA) performed independently within immune compartments reveals that donor-level scores along the first principal component (PC1) are strongly correlated across compartments at a fixed visit (Spearman correlation), demonstrating that shared donor ordering emerges prior to pooling across compartments. (B) Cross-compartment Spearman correlations of donor-level principal component scores are computed as a function of principal component index by projecting compartment-specific deviation profiles into a fixed reference PCA basis (defined in CD4 naive T cells). These correlations show that donor-level structure is shared across immune compartments and extends to additional principal components, supporting a multi-dimensional representation of donor-level variation. (C) Spearman correlation of donor-level PC1 scores between Year 1 Day 0 and Day 7 across matched donors shows strong preservation of donor ordering over short timescales (*ρ* = 0.86), indicating structured temporal coherence. (D) Comparison of PC1 scores between Day 0 and Day 90 shows substantially weaker correspondence (*ρ* = 0.30), indicating that donor-level transcriptional organization evolves over longer intervals. Together, these analyses establish that shared donor-level transcriptional structure is independently recoverable within compartments, extends across multiple principal components and exhibits temporally structured coherence.

To evaluate the biological coherence of this consolidated donor representation, we examine the temporal behavior of the first principal component score. For most donors, this score changes little between Year 1 Day 0 and Day 7 (Fig. 3C), but shifts more substantially between Day 0 and Day 90 (Fig. 3D), with the same pattern observed in Year 2. This indicates that the donor-level transcriptional organization captured by the first principal component is stable over short timescales yet evolves over longer ones. Temporal coherence is also observed across additional principal components, with characteristic timescales varying across components, supporting a structured biological organization rather than a single global complexity axis or purely technical variation.

These analyses show that coordinated transcriptional deviations across immune cell sub-types reflect a shared donor-level structure. This structure is captured by a multi-dimensional set of transcriptional axes, motivating a unified framework for representing donor-level transcriptional variation across immune cell types.

### A joint representation of donor-level transcriptional organization

To represent this shared structure in a unified framework, we apply PCA to deviation profiles pooled across subtypes and donor visits. The resulting principal components define a coordinate system in which donor-level transcriptional organization can be represented and quantified. For each principal component, we aggregate subtype-level scores across well-represented subtypes to obtain a single score for each donor–visit, such that each axis captures a distinct pattern of coordinated transcriptional variation across many genes. We refer to this unified representation as the immune transcriptional landscape (ITL).

Donor-level transcriptional variation lies in a low-dimensional space relative to the full gene space, with a small number of principal components capturing a substantial fraction of the total variance (Fig. 4A). This structure is broadly shared across immune cell types, as the same principal components account for a substantial fraction of variance within each subtype (Fig. 4B). However, the fraction of variance explained by individual components differs modestly across lineages, indicating that while donor-level structure is shared, its expression remains partially cell type-dependent. Importantly, each principal component reflects coordinated changes across many genes rather than a single transcriptional program, indicating that the ITL captures broad, distributed patterns of variation.

**Figure 4:**
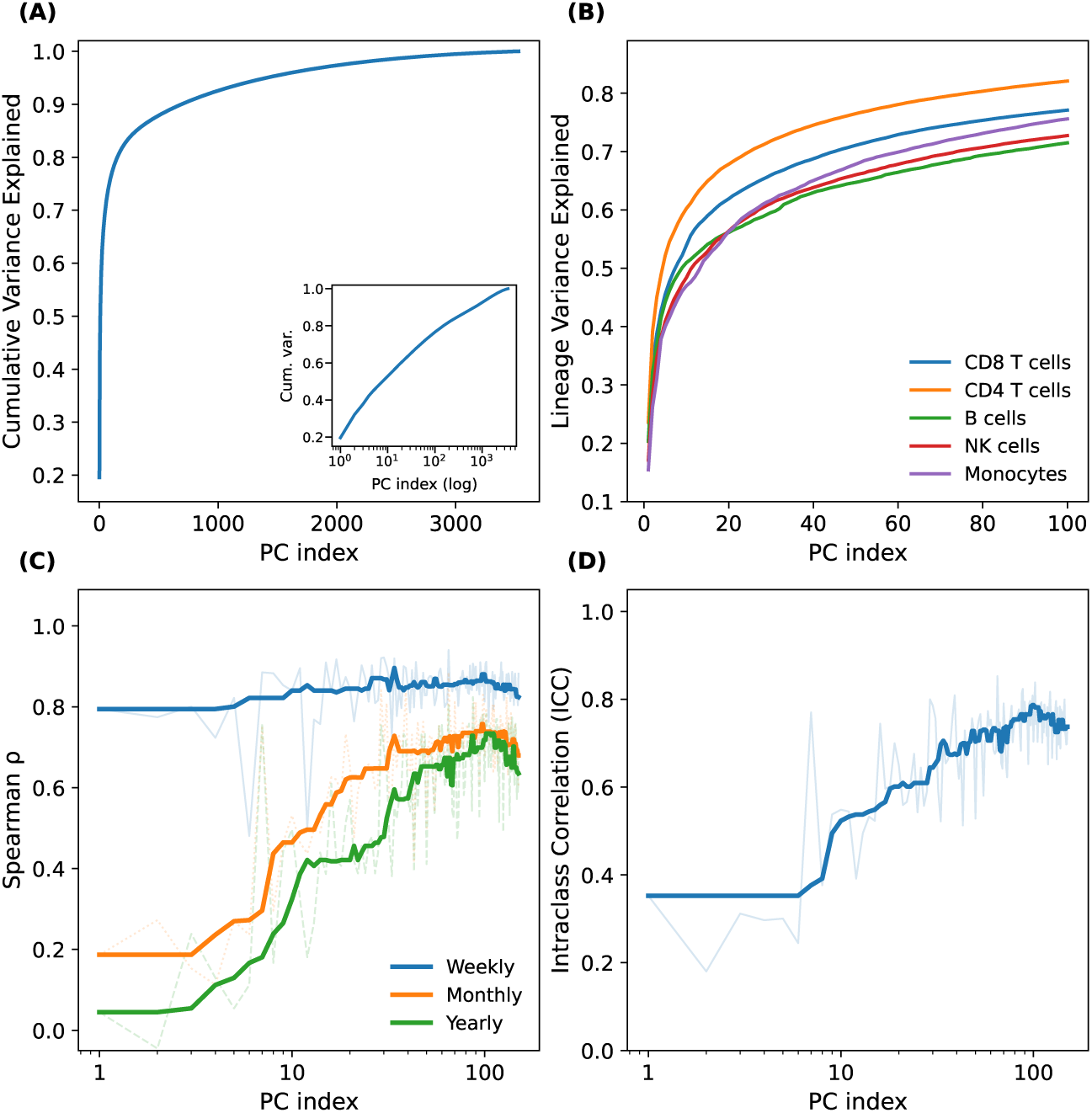
Shared and temporally structured donor-level immune organization. (A) Cumulative variance explained by principal component analysis (PCA) of pooled donor-level transcriptional deviation profiles, showing a compressed structure. Inset shows the same relationship on a logarithmic principal component index. (B) Fraction of within-lineage donor-level variance captured by the ITL axes, shown separately for CD8 T cells, CD4 T cells, B cells, NK cells and monocytes. A substantial portion of lineage-specific variance lies within the same shared multi-dimensional subspace. (C) Temporal coherence of donor ordering across ITL principal components, quantified as Spearman correlation of donor-level scores across visits at weekly, monthly and yearly intervals. Principal components differ in temporal persistence, with early components exhibiting only short-term coherence and later components retaining stability over longer timescales. Curves are averaged across annual cycles and a median smoothing is applied for clarity. (D) Donor-level stability of ITL principal components, quantified as intra-class correlation (ICC). Early components show greater within-donor variability, whereas many higher-order components exhibit stable donor-specific structure. Median smoothing is applied for clarity. Together, panels (C) and (D) demonstrate that the ITL resolves donor-level immune organization into shared principal components with distinct temporal and inter-donor stability profiles.

An important property of the ITL is that temporal coherence within donors varies systematically with component index (Fig. 4C). To quantify this, we compute the Spearman correlation of donor scores for each principal component across pairs of timepoints at different separations, such that high correlation indicates stability of donor scores, while low correlation reflects temporal reorganization. Leading components show high coherence over short intervals but decay over longer timescales, reflecting short-term transcriptional fluctuations shared across donors. In contrast, higher-order components exhibit lower short-term coherence but retain donor-specific variation over longer intervals. Intraclass correlation analysis further shows that stable donor-specific structure is distributed across multiple principal components (Fig. 4D), indicating that temporal and inter-individual variation are organized along distinct axes of the ITL. Notably, this stratified structure is not imposed by the analysis, as the ITL is constructed without incorporating temporal information, and therefore reflects underlying biological organization.

Although principal components are ordered by the variance they explain, biologically meaningful signals need not coincide with the largest sources of variation. Higher-order ITL components retain stable donor-specific structure (Fig. 4C), suggesting that they should support recovery of donor identity across time. To test this, we perform nearest-neighbor matching under a strict annual holdout, in which all Flu Year 2 samples are excluded during PCA fitting and subsequently projected into the learned basis. In this setting, PCs 100–150 recover donor identity across years with near-perfect accuracy, compared to 7% using the leading principal components (PCs 1–10), while permutation of donor labels reduces recovery to chance levels (mean accuracy 1%, empirical *p <* 0.001).

The ITL captures a donor-level transcriptional organization that is distributed across genes rather than confined to lineage-specific programs. As a consequence, even gene sets canonically associated with a specific lineage are expected to preserve donor-level ordering across other lineages. To test this, we use a canonical T cell receptor signaling gene set and, for each donor and visit, compute a program score as the mean deviation across these genes (see Methods). Importantly, these scores reflect deviations from lineage-specific baselines rather than absolute expression levels. Despite its canonical association with T cells, these deviation scores preserve donor-level ordering between memory T cells and other lineages, including naive T cells (*ρ* ≈ 0.96), B cells (*ρ* ≈ 0.97), NK cells (*ρ* ≈ 0.94) and myeloid populations (*ρ* ≈ 0.89). This result shows that donor-level transcriptional organization is shared across immune lineages even when evaluated using gene sets not canonically associated with those lineages. We observe the same pattern using a canonical B cell receptor signaling gene set, indicating that this coordination is not specific to any particular lineage-associated program but reflects a global donor-level structure.

To determine whether the temporal stratification observed in the ITL is directly reflected in gene expression, we evaluate temporal coherence for predefined gene sets. For each gene set, we aggregate deviation values across genes to obtain a donor-level score and quantify temporal coherence as the Spearman correlation of donor ordering across visits. Gene sets associated with early principal components (1–10) show strong short-term coherence but lose donor ordering over longer intervals (*ρ* ≈ 0.77 weekly vs. ≈ 0.16 yearly), whereas gene sets associated with later components (100–150) preserve donor ordering across yearly visits (*ρ* ≈ 0.63; see Methods). This pattern demonstrates that temporal stratification is directly observable in gene space and recapitulates the structure of the ITL (Fig. 4C). Importantly, distinct gene sets exhibit stability over different timescales, indicating that cross-lineage coordination cannot be explained by a single global complexity axis.

These observations indicate that immune transcription is organized not as independent lineage-specific programs or a single global axis, but as a coordinated, multi-dimensional donor-level structure with temporally stratified modes of variation. The cross-lineage shifts observed across immune compartments (Fig. 2) reflect this coordination in gene space, while the ITL resolves its structure into components, with leading components capturing longitudinal changes shared across donors and higher-order components capturing stable donor-specific differences. We next assess whether this organization generalizes across independent cohorts, tissues and measurement modalities.

### External replication and robustness of immune organization

To determine whether the donor-level transcriptional organization identified in the primary cohort reflects a general and robust property of human immune variation, we assess both external replication and stability to preprocessing and technical variation.

We first analyze an independent cross-sectional PBMC cohort of 166 healthy donors generated using a distinct study design and processing pipeline [16]. Applying the same analytical framework, we observe that deviation profiles correlate across immune lineages within donors (Fig. 5A), PCA recovers a shared donor ordering across compartments (Fig. 5B), and cross-compartment coherence extends across multiple principal components (Fig. 5C). Because the external cohort is cross-sectional, these analyses assess replication of cross-lineage deviation alignment and shared donor-level structure but do not test the temporal stratification observed in the longitudinal Sound Life cohort.

**Figure 5:**
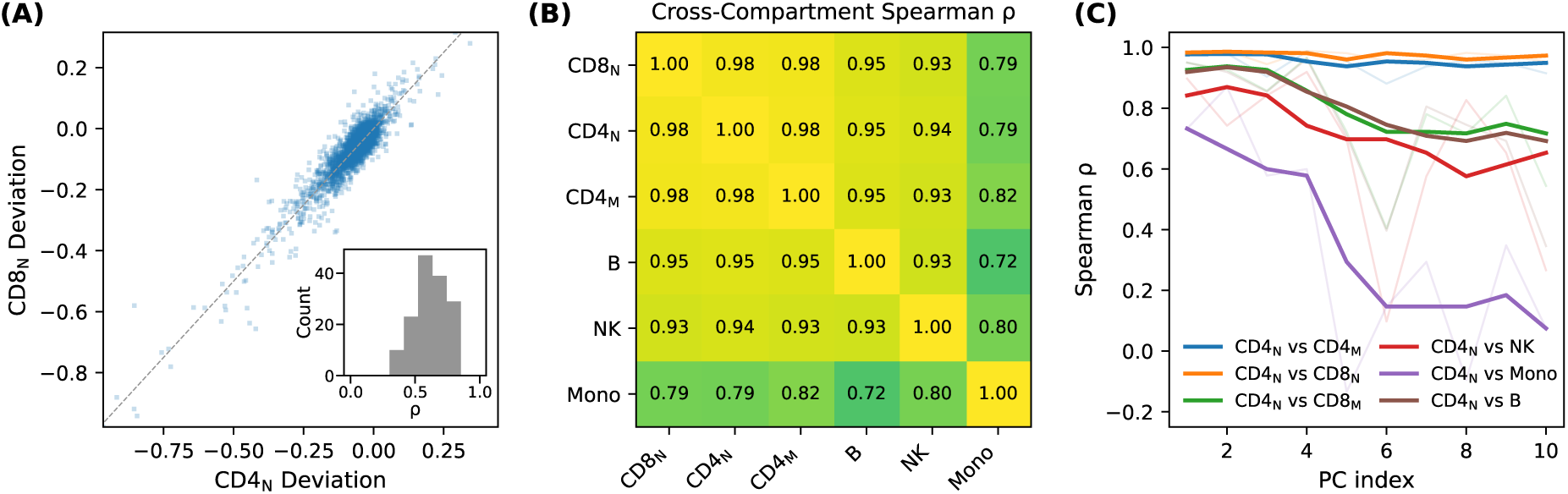
Independent replication of shared donor-level transcriptional structure in an external cohort. (A) Direct comparison of gene-wise transcriptional deviations between immune cell types in an independent healthy PBMC cohort shows positive correlation after centering relative to cell type-specific reference profiles, indicating coordinated deviations across cell types without dimensionality reduction. (B) Spearman correlation matrix of donor ordering along the first principal component of a joint PCA fit to the external cohort, with PC1 aggregated within immune compartments, demonstrates strong cross-compartment agreement of the primary donor-level component. (C) Cross-lineage correspondence of donor ordering as a function of principal component index, evaluated using representative anchor subtypes for each lineage, shows that shared donor-level structure extends beyond the first principal component to additional principal components. Together, these analyses demonstrate that the shared donor-level transcriptional structure motivating construction of the immune transcriptional landscape is reproducible in an independent cohort, supporting the robustness and generality of the ITL framework.

We next evaluate whether this structure is robust to preprocessing choices and technical variation. The overall geometry and temporal coherence of the donor-level structure remain stable under multiple perturbations of the analysis. These include fitting PCA after excluding Flu Year 2 Day 0 and Day 7 samples, recomputing subtype centering using training data only and projecting held-out data into the learned basis, and downsampling the number of cells contributing to each donor–visit–subtype profile prior to aggregation. We also evaluate alternative aggregation strategies, including averaging per-cell normalized expression values and pseudobulk aggregation based on summed counts, both of which yield consistent results. The ITL is similarly stable under alternative gene prevalence thresholds and exclusion of activation-associated or high-loading genes. Although the first principal component correlates with mean detected genes and total UMI counts, consistent with known relationships between transcriptional output and sequencing complexity [24], this association is largely confined to PC1. Because sequencing complexity can reflect both technical capture efficiency and biological differences in transcriptional output, we do not treat this variation as purely technical or remove it as a nuisance effect. The donor-level immune organization described here is distributed across many axes, remains evident beyond the first principal component and exhibits temporal stratification, indicating that it is not explained by sequencing complexity.

These analyses indicate that the donor-level structure captured by the ITL is not reducible to preprocessing choices, sampling variability or simple sequencing-related effects.

### Immune organization extends across tissues and protein-level measurements

We next examine whether the coordinated donor-level transcriptional variation identified in peripheral blood generalizes across tissues and measurement modalities. To test this, we analyze a multimodal single-cell dataset comprising paired RNA and surface protein (CITE-seq) measurements from over one million immune cells profiled across blood, lymphoid and mucosal tissues from 24 deceased human organ donors [17]. Analyses are restricted to tissue and cell type combinations with sufficient donor overlap and are interpreted as assessing generalization across a diverse but heterogeneously sampled set of immune contexts. Deviation profiles are computed as in the Sound Life analysis by centering expression within each cell type and tissue, isolating donor-specific variation from baseline tissue differences. We examine donor-matched deviation correlations across cell types within tissues and across tissues within cell types using both RNA and CITE-seq protein measurements (see Methods).

Within tissues, deviation profiles from different immune cell types are positively correlated within individual donors in RNA (Fig. 6A). Across tissues, deviation profiles for the same cell type are likewise positively correlated within donors (Fig. 6B). Similarly, CITE-seq measurements show coordinated variation both across immune cell types within tissues and across tissues within cell types (Fig. 6C,D), mirroring the transcriptional alignment observed in RNA. In all cases, these correlations are lost under donor permutation, confirming that the observed structure depends on within-donor correspondence.

**Figure 6:**
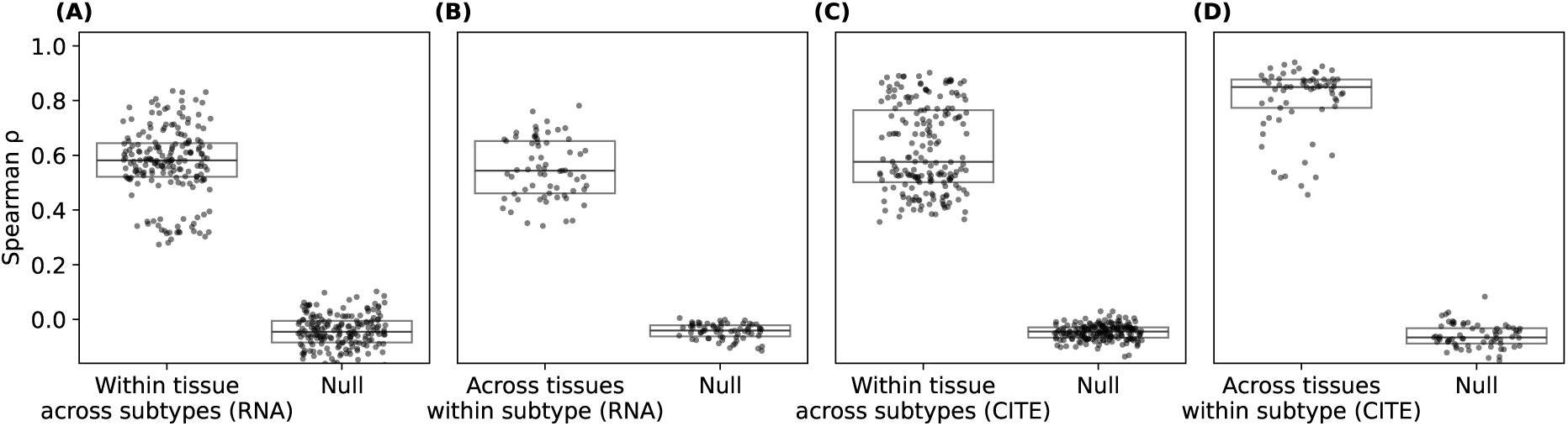
Coordinated donor-level immune variation across tissues and molecular modalities. Donor-matched deviation correlations were computed across cell types within tissues and across tissues within cell types using both RNA and CITE-seq protein measurements. In all panels, deviation profiles were centered within each comparison group and correlations were computed across matched donors, with a null distribution generated by donor permutation within matched sets. (A) Within tissues, deviation profiles from different immune cell types are positively correlated across donors in RNA, indicating coordinated transcriptional variation across cell types within a shared tissue environment. (B) Across tissues, deviation profiles for the same cell type remain positively correlated within donors in RNA, demonstrating that donor-specific transcriptional state is preserved across tissues. (C) Within tissues, analogous analyses using CITE-seq protein measurements show coordinated variation across immune cell types at the protein level, indicating that cross-cell-type coordination is reflected in cellular phenotype. (D) Across tissues, protein-level deviation profiles for the same cell type are strongly correlated within donors, demonstrating preservation of donor-specific immune state across tissues at the level of protein expression. Together, these analyses show that coordinated donor-level immune organization is not restricted to a single tissue or transcriptional measurements, but extends across multiple tissues and is reflected in both gene expression and protein-level phenotypes.

These results show that donor-level variation is coordinated across cell types within tissues, maintained across anatomical compartments for individual cell types and reflected at both the transcriptional and protein levels. These patterns are consistent with coordinated donor-level immune variation spanning both peripheral and tissue-resident immune cell populations.

### Immune organization across circulating proteomics and physiological measurements

We next examine whether the donor-level structure captured by the ITL is reflected in independent circulating measurements, including routine hematologic indices and serum proteomic profiles measured across a broad panel of circulating proteins. Donor positions within the ITL occupy a constrained region of this high-dimensional space. Thus, for these analyses we summarize donor positions in ITL space by applying PCA to the first 150 ITL components. This threshold is motivated by temporal coherence analysis and the general observation that higher-order principal components increasingly capture noise. Components within this range retain structured, longitudinally coherent signal (Fig. 4C), indicating that they remain well within the regime of biologically meaningful variation. Whereas the ITL defines axes of coordinated transcriptional variation across genes, this reduced representation captures the dominant axes of variation in donor immune state. For proteomic measurements, we derive an analogous low-dimensional representation, enabling direct comparison of donor positions across transcriptional and proteomic spaces (see Methods).

Correlation of components of the reduced ITL representation with routine hematologic indices reveals a structured pattern. The first few components show minimal association (*ρ* ≈ 0.0 − 0.1), intermediate components show modest correlations (*ρ* ∼ 0.1 − 0.25), and later components exhibit stronger associations, most notably with hemoglobin and hematocrit (*ρ* ∼ 0.4, *p <* 10^−30^) and to a lesser extent platelet count (*ρ* ∼ 0.25, *p <* 10^−15^). Thus, classical blood physiology is reflected in identifiable directions of the reduced ITL but does not account for its dominant transcriptional structure.

Extending this cross-modal evaluation to molecular measurements, we next assess correspondence between donor-level structure in the ITL and circulating proteomic space. Using the same reduced ITL representation, we compare donor positions across transcriptional and proteomic manifolds. For a single illustrative ITL axis, donor ordering defined by transcriptional variation is accurately recovered from proteomic features in held-out data (Fig. 7A), demonstrating alignment between modalities. To evaluate correspondence beyond a single axis, we compare donor positions across the first ten principal components of both representations under visit-level and donor-level holdout. In each case, cross-modal alignment exceeds permutation expectations (Fig. 7B-C). These results indicate that donor-level immune organization is coherently reflected in circulating proteomic measurements.

**Figure 7:**
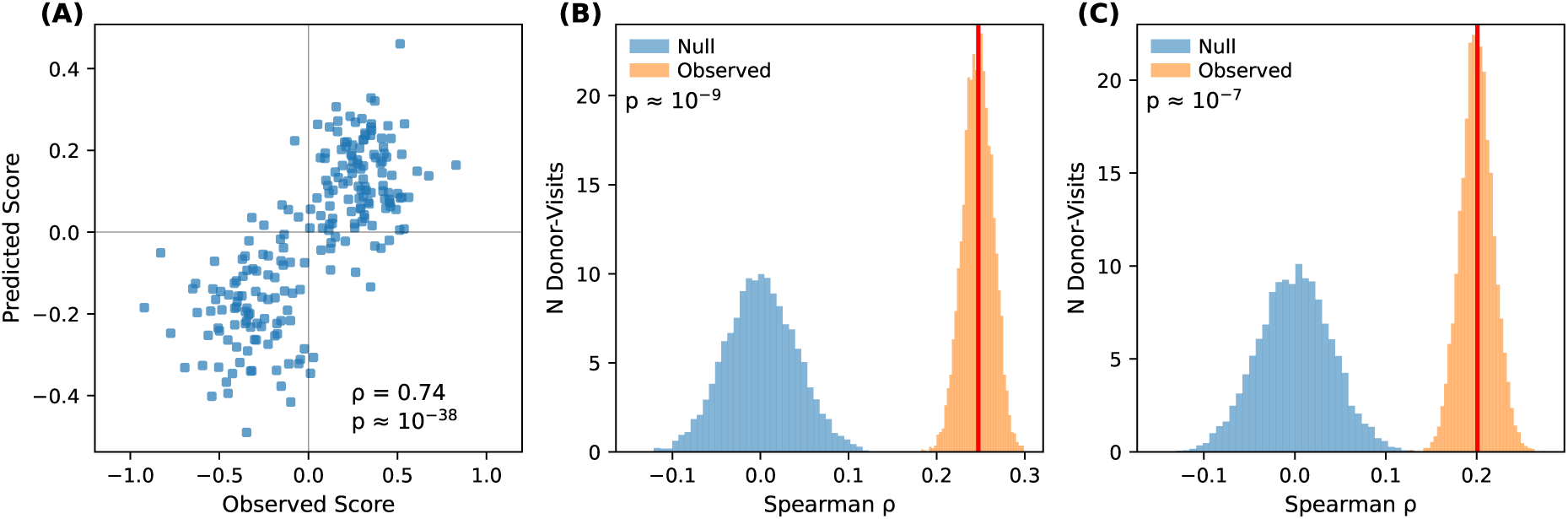
Correspondence between immune transcriptional landscape position and circulating serum proteomics. (A) Out-of-sample correspondence between observed donor-level score along one principal component (PC7) of the reduced ITL and values predicted from serum proteomic features. Proteomic features are mapped to ITL scores using ridge regression with donor holdout and correspondence is quantified by Spearman correlation. (B) Multivariate assessment of proteomics-ITL correspondence under temporal holdout, simultaneously modeling the first ten principal components of the reduced ITL representation. Proteomic features are mapped to these axes using multi-output ridge regression and correspondence is quantified as the mean Spearman correlation across axes on held-out visits. The observed distribution (bootstrap resampling of test samples) is compared to a permutation null in which the correspondence between proteomic features and ITL scores is disrupted by randomly reassigning ITL targets across donors in the training set. (C) Corresponding robustness analysis under donor holdout, in which entire donors are withheld during training. The observed correspondence again exceeds the permutation null, demonstrating that the proteomics-ITL relationship generalizes to unseen donors. Together, these analyses show that donor-level position within the immune transcriptional landscape is reflected in independent circulating protein profiles.

Coordinated transcriptional variation across immune cell types is temporally stratified and reflects a donor-level organization that is reproducible across cohorts, extends across tissues and is independently captured by cellular phenotype and circulating measurements. Notably, this structure is not readily explained by shared sample-level technical effects and instead supports a biological origin of the observed coordination consistent with a shared donor-level immune state. We next assess whether this organization is preserved under immune perturbation.

### Donor-level transcriptional organization is preserved under perturbation

The immune system is continuously exposed to endogenous and exogenous perturbations. If perturbations completely overwrite donor-level transcriptional organization, persistent coordination would not be expected. Instead, the observation of stable donor-level organization in the Sound Life cohort suggests that donor-level transcriptional coordination is preserved under perturbation. To test this hypothesis, we analyze perturbation responses to determine whether they remain coordinated across immune cell types within individuals.

The Parse Biosciences dataset [18] profiles PBMCs from 12 donors stimulated *in vitro* with 90 cytokines. To enable direct comparison with the Sound Life cohort, we aggregate data to donor-level mean expression profiles for each cell type and perturbation. Responses are defined relative to donor-matched unperturbed baselines, yielding deviation profiles that place perturbation responses in the same relative expression space used to define the ITL.

We restrict analyses to a shared gene set and project perturbation responses into the ITL defined from Sound Life. Under this matched representation, a substantial fraction of variance in perturbation responses is captured within the same low-dimensional space that describes natural variation (Fig. 8A), indicating that in vitro perturbation responses are preferentially aligned with this space rather than inducing orthogonal states. Within-donor coordination of transcriptional responses across cell types remains strong across perturbations (Fig. 8B). To distinguish donor-specific coordination from coordinated responses that are shared across cell types, we compare the observed coordination with a donor-permuted control in which donor matching is disrupted while preserving coordination arising from shared responses to each cytokine. Across all 90 cytokines, within-donor coordination exceeds the donor-permuted null, demonstrating that donor-specific coordination is present for every cytokine examined.

**Figure 8:**
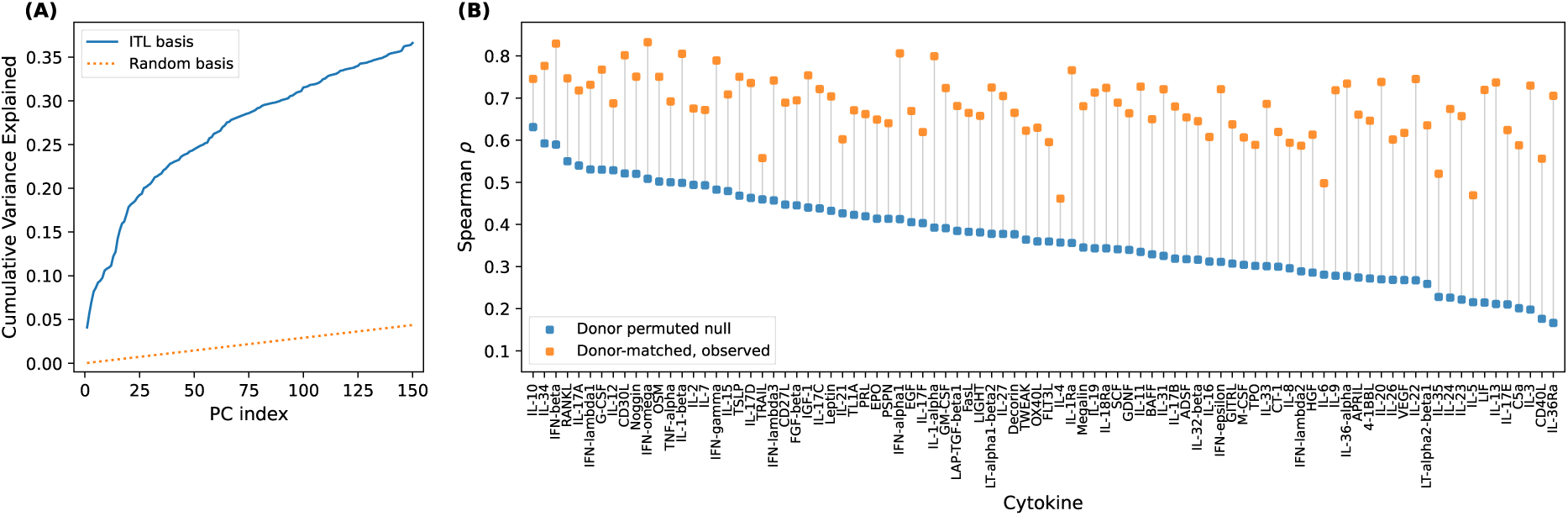
Perturbation responses align with the immune transcriptional landscape and remain coordinated across immune cell types. (A) Cumulative variance of the Parse dataset projected into the ITL basis (blue) compared to a random orthonormal basis (dashed), showing that perturbation responses are preferentially aligned with the space defined by natural variation. (B) Cross-cell-type coordination of perturbation responses for all ninety cytokines. For each cytokine, within-donor coordination was quantified as the median pairwise Spearman correlation across cell type-specific perturbation response vectors. Cytokines are ordered by the median coordination observed after donor permutation, which preserves coordination arising from shared responses to each cytokine while disrupting donor matching across cell types. Observed coordination (median *ρ* = 0.68) exceeds the donor-permuted null (median *ρ* = 0.36) for every cytokine examined, demonstrating a donor-specific component of coordinated perturbation responses across immune cell types.

Perturbation responses preserve donor-level transcriptional coordination across immune cell types, supporting the expectation that perturbations do not overwrite the donor-level organization observed in the Sound Life cohort. Together with the longitudinal analyses of healthy donors, these findings suggest that coordinated transcriptional organization may be a fundamental organizing principle of immune state, characterizing both donor transcriptional state and transcriptional responses to perturbation.

## Discussion

Immune organization is reflected in coordinated transcriptional variation across cell types within individuals, defining a low-dimensional state space that captures longitudinal change and baseline differences between individuals. Importantly, this coordinated organization is evident not only in transcriptional state but also in transcriptional responses to perturbation, indicating that transcriptional coordination characterizes both immune state and immune response. Within this space, dominant patterns reflect longitudinal dynamics shared across individuals, whereas others encode stable individual-specific structure that persists across annual visits and enables recovery of donor identity from expression profiles. Despite the complexity of the immune system, these results point to a simple underlying principle: variation in gene expression takes the form of coordinated, donor-specific deviations shared across cell types, such that even canonical lineage-associated programs exhibit coherent shifts across immune compartments rather than varying independently within each cell type. This organization extends across tissues and measurement modalities, indicating that cellular responses in the immune system do not vary independently, but instead reflect coordinated manifestations of an individual-specific state.

The immune system operates as an integrated network of diverse cell types [25], yet immune organization is typically characterized through cell-type composition, signaling interactions and circulating measurements. Because transcription integrates upstream regulatory inputs and shapes downstream cellular behavior [26, 27], it provides a natural layer at which system-level immune organization can be observed. The immune transcriptional landscape (ITL) defines a coordinate system for this structure, enabling quantification across individuals and over time. Importantly, this organization is not specific to transcription alone: the coordinated structure we identify extends across tissues and is independently reflected in cellular phenotype and circulating measurements, including routine hematologic indices and matched proteomic profiles [23, 28]. These observations indicate that the ITL provides a coordinate system for representing an underlying individual-level immune state.

Our findings are consistent with prior observations that the human immune system exhibits substantial inter-individual diversity while remaining relatively stable within individuals over time [20, 21, 22, 23]. Persistent immune signatures and individual-specific set points have been described across multiple cohorts [29, 30, 12]. Here we resolve these phenomena at the level of transcription, showing that stability and diversity are embedded within a shared transcriptional state space. This individual-level structure aligns with observations across immune compartments [8, 25, 3], and the ITL provides a quantitative framework for representing and analyzing this immune state directly in transcriptional space.

The individual-level organization revealed by our findings constrains plausible mechanisms of immune regulation. Transcriptional coordination across compartments indicates that dominant drivers act in parallel across cell types rather than being cell type-intrinsic [31, 32]. This structure spans thousands of genes and persists after exclusion of canonical activation-associated programs, consistent with distributed, transcriptome-wide modulation rather than regulation by a limited set of pathways [33, 34]. The coexistence of rapidly varying and stable patterns further suggests regulatory processes operating across multiple timescales [35, 33, 22]. Together with the observation of this structure across tissues and measurement modalities, these features point to systemic regulators that couple immune compartments, including physiological and environmental influences, circulating cytokine and protein signaling and stable individual-specific regulatory architecture shaped by genetic and non-heritable factors [35, 33, 34, 23, 28, 20, 36, 31].

We characterize individual-level immune organization in healthy adults. An important next step is to determine how defined immune perturbations, such as infection, vaccination and physiological or environmental changes, reshape trajectories in the transcriptional state space, and how responses depend on baseline immune context, including age and sex. In the present study, we aggregate expression across many cells within each subtype and therefore do not resolve fine-grained cell-to-cell heterogeneity. Single-cell analyses could reveal how this coordinated transcriptional variation arises at the level of individual cells. While this coordinated structure is evident, the upstream regulatory processes underlying it remain unspecified. Extending this framework to integrative multiomic analyses will help identify the causal drivers of immune organization and its alteration in disease.

We find that coordinated transcriptional variation across immune cell types within individuals reflects a shared, system-level organization of the immune system. This organization links variation in gene expression across cell types to independent systemic measurements and is evident in both transcriptional state and transcriptional responses to perturbation. These results provide a foundation for investigating how individual-specific set points are maintained, how regulatory processes operate across cell types and how coordinated transcriptional organization is reshaped across aging, environmental exposure, physiological perturbation and disease.

## Methods

### Cohorts

#### Sound Life longitudinal cohort

Primary analyses use data from the Sound Life longitudinal healthy adult cohort, a prospective multiomic study with repeated immune profiling of the same individuals over time [19]. Participants are sampled across annual cycles with a standardized visit structure including baseline (Day 0), short-term (Day 7) and intermediate-term (Day 90) follow-up visits, with seasonal influenza vaccination administered at Day 0 of each cycle. In addition to these core visits, a subset of donors contributes additional intermediate visits. The same individuals are followed across multiple years, enabling within-subject analysis of immune transcriptional organization over weeks, months and years.

#### Independent healthy PBMC atlas

To assess generality of the transcriptional structure identified in the Sound Life cohort, we analyze an independent large-scale single-cell atlas of healthy human peripheral blood mononuclear cells described by Terekhova et al. [16]. This cohort comprises single-cell transcriptomic profiles from 166 healthy donors and represents a primarily cross-sectional design. Raw UMI count matrices and subtype annotations are obtained and processed using the same normalization, filtering and aggregation workflow applied to the Sound Life data, as described below. Analyses in the external cohort are restricted to the Sound Life ITL gene universe, retaining the intersection of genes present in both datasets (all but one gene). The external cohort is used exclusively to test whether cross-lineage deviation alignment and shared donor-level structure are present in an independent dataset; no projection of external samples into the Sound Life ITL is performed.

#### Cross-tissue immune atlas

To assess generalization of donor-level transcriptional organization across anatomical contexts and molecular modalities, we analyze a multimodal single-cell dataset of human immune cells profiled across tissues by Wells et al. [17]. This cohort comprises paired RNA and surface protein (CITE-seq) measurements from over one million immune cells isolated from blood, lymphoid and mucosal tissues obtained from 24 deceased human organ donors. Tissue sampling includes multiple immune-relevant sites, such as blood, bone marrow, spleen, lymph nodes, lung and intestinal compartments, although tissue availability and cell numbers vary across donors. Data are processed using the same normalization, filtering and aggregation framework applied to the Sound Life cohort. Analyses are restricted to cell types shared with the ITL framework and to tissue and cell type combinations with sufficient donor overlap. This dataset is used exclusively to evaluate whether coordinated donor-level variation is preserved across tissues and reflected at the level of surface protein expression.

#### Parse cytokine perturbation dataset

To examine how donor-level transcriptional organization relates to perturbation responses, we analyze a large-scale single-cell cytokine perturbation dataset generated by Parse Biosciences [18]. This dataset comprises peripheral blood mononuclear cells (PBMCs) from 12 healthy donors stimulated in vitro for 24 hours with 90 individual cytokines or PBS control, with transcriptional profiles measured using combinatorial barcoding. Cytokines were selected to span major immune signaling pathways, including interferon, interleukin and growth factor families, providing a comprehensive panel of physiologically relevant immune perturbations. The dataset is used to assess how perturbation-induced transcriptional responses relate to the donor-level organization identified in the Sound Life cohort.

#### Starting data and preprocessing state

For all datasets, analyses begin from post-quality-control, annotated single-cell data as provided by the original studies. Cell-level quality control, doublet removal and initial cell type annotation are performed by the original study authors [19, 16, 17, 18] and are not repeated here. For the cross-tissue immune atlas, we use the processed AnnData object provided with the study, extract donor, tissue and native cell type annotations, restrict RNA features to genes overlapping the ITL gene universe and aggregate expression to donor by tissue by cell type means. CITE-seq protein measurements are taken from the normalized CITE matrix provided in the object and aggregated over the same retained cells and groups. Downstream analyses then apply analysis-specific restrictions, including cell type selection, minimum cell counts per group and donor overlap requirements, as described below.

#### Cell filtering, normalization and aggregation

Analyses operate on post-quality-control, annotated single-cell datasets as provided by the original studies. Where raw count matrices are available, gene expression values are library-size normalized to a common target sum *T* = 4524 and transformed using log(1 + *x*). Specifically, if *x_ig_*denotes the count for gene *g* in cell *i*, and *L_i_* = Lg *x_ig_*is the library size of cell *i*, we compute

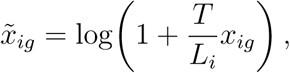

for cells with *L_i_ >* 0.

To reduce the influence of transient non-homeostatic states on donor-level baseline structure, cells exhibiting strong transient activation, cell-cycle or interferon-associated signatures are filtered using subtype-specific thresholds computed within L3 labels. For each cell, module scores for three predefined gene sets (cell-cycle S, cell-cycle G2M, and a core interferon-stimulated gene set) are computed as the mean of *x^-^_ig_* over genes in the set. For each L3 subtype, a cutoff is defined as the 99th percentile of the corresponding score distribution across retained cells, and cells are excluded if they exceed the subtype-specific cutoff for any of the three scores. Results are insensitive to the exact cutoff used.

For each donor–visit–L3 group, subtype-resolved mean expression profiles are computed by taking the arithmetic mean of *x^-^_ig_* across all retained cells in the group for each gene. Only groups containing at least 100 cells are retained for downstream analysis. The resulting group-by-gene matrix of mean log-normalized expression values forms the basis for gene filtering, reference profile centering and dimensionality reduction.

#### Gene universe definition

To define a consistent gene universe for dimensionality reduction and cross cell type comparison, we first performed gene filtering separately within each Sound Life compartment or lineage pipeline. Gene prevalence was defined as the fraction of retained cells with nonzero raw counts for a given gene and was computed within coarse label groups for T cell compartments or within L3 subtype labels for non T lineages. Within each pipeline, genes were retained if prevalence exceeded 10% in every label group, after excluding mitochondrial and ribosomal genes. The final immune wide ITL gene universe was defined as the intersection of retained genes across the CD4 naive, CD4 memory, CD8 naive, CD8 memory, B cell, NK cell and monocyte pipelines, and this fixed Sound Life derived gene set was used for downstream analyses and external datasets.

Mitochondrial and ribosomal genes are excluded from the retained gene universe prior to principal component analysis. No additional exclusion of activation-associated genes was applied in the primary gene universe. Instead, transient non-homeostatic states were controlled at the cell level through the filtering procedure described above. Exclusion of selected activation-related gene programs was performed only in dedicated sensitivity analyses. Genes with names beginning with the prefix `MT-` are classified as mitochondrial genes, and genes with prefixes `RPL` or `RPS` are classified as ribosomal genes. Although excluded from the PCA gene set, aggregate mitochondrial and ribosomal expression levels are computed separately for each donor–visit–subtype group and retained for diagnostic and alignment analyses.

#### Reference profile centering and deviation profiles

Subtype-resolved mean expression profiles are centered to define subtype-specific reference profiles. For each L3 subtype, the reference profile expression vector is defined as the mean of subtype-resolved group-level expression profiles across all donors and visits retained for analysis. Deviation profiles are computed by subtracting this subtype reference profile from each donor–visit–subtype mean expression profile. These centered deviation profiles remove systematic subtype-specific expression differences while preserving coordinated variation across donors and visits, and serve as input to deviation alignment analyses and principal component analysis. Except where noted for illustrative purposes (Fig. 2A), all downstream analyses operate on these donor–visit–subtype deviation profiles or linear transformations derived from them.

#### Deviation alignment analyses

To assess whether transcriptional deviations from cell type-specific reference profiles are shared across immune cell types within individuals, we analyze centered deviation profiles defined as described above.

#### Illustrative single donor–visit analysis

For visualization, a representative donor–visit with high cell counts in paired immune subtypes is selected. Gene-wise Spearman correlation across deviation profiles is used to quantify alignment between cell types, illustrating the effect of subtype reference profile centering on cross-cell type correspondence.

#### Population-level deviation alignment

For population-level analysis, gene-wise Spearman correlations are computed between deviation profiles for all donor–visit combinations in which the relevant cell type pairs are present. Correlations are calculated across genes independently for each donor–visit, yielding a distribution of correlation coefficients across the cohort. This procedure is applied uniformly to T cell subsets, B cells, NK cells and monocytes using the same centering strategy.

#### Null model construction

Statistical significance of deviation alignment is evaluated using a visit-preserving donor-shuffled null model. For each visit independently, donor identities are permuted with the constraint that no donor is paired with itself. Deviation profiles from one cell type are paired with deviation profiles from a different donor at the same visit, preserving visit-specific context while disrupting within-individual correspondence. The resulting distribution of gene-wise Spearman correlations defines the null expectation.

#### Compartment-specific PCA and cross-compartment coherence

To characterize donor-level transcriptional structure within immune compartments, principal component analysis (PCA) is performed independently within each major compartment using deviation profiles as input. PCA is fit using randomized singular value decomposition with a fixed random seed. Principal component scores and gene loading vectors are retained for downstream analyses.

#### Donor-level aggregation

For cross-compartment comparisons, subtype-level principal component scores are aggregated to donor–visit level by averaging across L3 subtypes and then averaging across coarse compartment labels (L2) within each donor at a fixed visit (L3 → L2 → donor). This produces a single donor-level score per principal component for each compartment and visit.

#### Cross-compartment coherence of the first principal component

To assess whether dominant donor-level structure is shared across immune compartments, donor-level scores for the first principal component are compared across compartments at a fixed visit using Spearman correlation, yielding a cross-compartment correlation matrix.

#### Cross-compartment coherence across principal components

To evaluate whether shared donor-level structure extends beyond the first principal component, deviation profiles from each target compartment are projected into a fixed PCA basis defined in a reference compartment. Let *X*^(^*^t^*^)^ ∈ R*^nt^*^×^*^G^* denote the centered group-by-gene matrix for target compartment *t*, and let *L*^(^*^r^*^)^ ∈ R*^k^*^×^*^G^* denote the loading vectors corresponding to the first *k* principal components from the reference compartment, with all compartments restricted to a shared ordered gene list. Projected scores are computed as

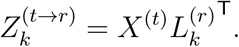

Projected scores are aggregated to donor–visit level as described above. For each principal component index *j* ∈ {1*, …, k*}, donor-level scores from the reference compartment (native PCA space) are compared to donor-level projected scores from each target compartment using Spearman correlation at a fixed visit. Correlation curves as a function of principal component index summarize cross-compartment coherence across leading and higher-order principal components.

#### Construction of the immune transcriptional landscape

The immune transcriptional landscape (ITL) is constructed by pooling donor–visit–subtype deviation profiles across immune compartments and performing principal component analysis (PCA) on the resulting joint matrix. PCA is fit with `n components=None`, retaining all principal components of the joint matrix. For each input pipeline (cd8 naive, cd8 memory, cd4 naive, cd4 memory, b, nk, mono), a shared gene list is defined as the intersection of `genes kept` across all pipelines and each pipeline’s group-level mean expression matrix (`group means`) is subset and reindexed to this shared ordered gene list. The resulting matrices are concatenated row-wise to form a joint group-by-gene matrix of deviation profiles. In total, this matrix contains 14,979 deviation profiles across 3,530 genes.

Principal component analysis is then performed on this joint matrix using randomized singular value decomposition with a fixed random seed. Principal component scores, gene loading vectors and variance explained are retained. The resulting principal components define the ITL axes.

#### Variance captured within lineages by the ITL subspace

To quantify how much within-lineage variance is captured by the ITL subspace (Fig. 4B), we use the joint centered matrix and the ITL gene-space basis. Let *X_L_*denote the subset of rows of the joint centered matrix corresponding to lineage *L* (mapping sources to five lineages: CD8 includes cd8 naive and cd8 memory; CD4 includes cd4 naive and cd4 memory; B, NK, and Mono correspond to b, nk, and mono). Let *U_k_* denote the first *k* ITL axes in gene space (rows of `pca model.components`). For each lineage, the fraction of variance captured by the first *k* ITL axes is computed as

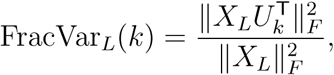

where ‖‧‖*_F_* denotes the Frobenius norm. Curves FracVar*_L_*(*k*) are evaluated over component index *k* using a fixed maximum *K*_max_.

#### Temporal coherence across ITL axes

Temporal coherence of donor-level ITL structure across axes (Fig. 4C) is evaluated using donor–visit ITL coordinates constructed by robust aggregation across high-prevalence L3 subtypes. An L3 subtype is considered present for a donor if it appears in the ITL `pca scores` for that donor at any visit. Subtypes present in at least 95% of donors are retained, and analyses are restricted to the six canonical visits (Flu Year 1 Day 0, Day 7, Day 90; Flu Year 2 Day 0, Day 7, Day 90). Donor–visit coordinates are computed by averaging ITL principal component scores across the retained high-prevalence L3 subtypes (direct L3 → donor–visit aggregation without intermediate L2 averaging).

For each pair of visits (*v*_1_*, v*_2_), temporal coherence is quantified for each ITL axis by computing the Spearman correlation of donor-level scores between *v*_1_ and *v*_2_ across donors with measurements at both visits. Short-term coherence is computed for Day 0 vs Day 7 within each year and averaged across Year 1 and Year 2; intermediate-term coherence is computed for Day 0 vs Day 90 within each year and averaged across Year 1 and Year 2; and long-term coherence is computed using matched cross-year comparisons (Year 1 Day 0 vs Year 2 Day 0 and Year 1 Day 7 vs Year 2 Day 7), averaged across the two comparisons. For visualization, correlation curves across axis index are smoothed using a running median with a fixed window size (window size = 11).

#### Intraclass correlation across ITL axes

To quantify the extent to which each ITL axis reflects stable between-donor differences versus within-donor fluctuations (Fig. 4D), we compute an intraclass correlation-like statistic on the same donor–visit ITL coordinates used for temporal coherence. Using the robust L3 subtype set (95% donor prevalence) and the six canonical visits, donor–visit scores are obtained by averaging ITL principal component scores across retained L3 subtypes.

For each ITL axis *i* (restricted to the first 150 axes), between-donor variance is computed as the variance across donors of the donor mean score averaged across visits, Var*_d_*(E*_v_*[PC*_i_*]). Within-donor variance is computed as the mean across donors of the variance across visits within each donor, E*_d_*[Var*_v_*(PC*_i_*)]. The intraclass correlation statistic is defined as

High values indicate axes dominated by stable donor-specific differences, whereas low values indicate axes dominated by within-donor temporal variation.

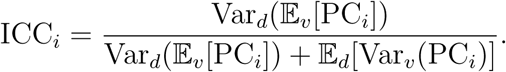

#### Gene program coherence and temporal stratification analysis

To evaluate how predefined gene programs reflect donor-level transcriptional variation, we operate directly on deviation profiles computed as described above. Gene program scores are computed by averaging deviation values across genes in a given set, yielding a scalar score per donor × visit × subtype without projection onto principal components.

To assess cross-lineage coherence, canonical T cell and B cell receptor signaling gene sets were obtained from the KEGG pathway database [37, 38]. For each canonical visit, program scores are aggregated within major immune compartments (T memory, T naive, B cell, NK cell, and myeloid) and Spearman correlations are computed across donors between compartments. Temporal coherence is assessed by averaging program scores across well-sampled compartments to obtain a donor-level score per visit, followed by pairwise Spearman correlations across visits.

To examine temporal structure independent of PCA, gene sets are also defined using ITL loadings by selecting the top 75 genes with highest mean absolute loading across either early (PC1–10; dynamic) or late (PC100–150; stable) components. These gene sets are scored identically in deviation space and temporal coherence is evaluated across predefined timescales of the canonical visits (weekly: Day 0–7; monthly: Day 0–90; yearly: matched visits across seasons).

#### External cohort analyses

Analyses in the independent healthy PBMC atlas [16] test whether the empirical conditions motivating construction of the immune transcriptional landscape in the Sound Life cohort are present in an independent dataset. Raw UMI count matrices from the external cohort are processed using the same preprocessing and analysis pipeline described above, including normalization, cell filtering, aggregation to subtype-resolved mean expression profiles, gene universe restriction and reference profile centering.

All analyses therefore operate on deviation profiles constructed using the identical workflow applied to the Sound Life cohort. External samples are not projected into the Sound Life ITL, and no model parameters or representations are transferred between cohorts.

To evaluate cross-lineage transcriptional alignment in the external cohort, gene-wise Spearman correlations are computed between deviation profiles from representative lineage pairs within individual donors, mirroring the deviation alignment analyses performed in the Sound Life cohort.

To assess donor-level transcriptional structure, principal component analysis is performed on centered subtype-resolved mean expression profiles using the same procedure applied in the Sound Life analyses. Donor-level scores for the first principal component are aggregated within compartments by averaging across L3 subtypes and compared across compartments at a fixed visit using Spearman correlation.

Finally, correlations across donors are evaluated as a function of principal component index to determine whether multi-dimensional donor-level transcriptional organization, as observed in the Sound Life cohort, is also present in the independent dataset.

#### Robustness and Technical Sensitivity Analyses

To evaluate the robustness of the Immune Transcriptional Landscape (ITL) to technical and preprocessing choices, we performed a series of targeted sensitivity analyses designed to test whether the observed donor-level transcriptional structure depends on specific modeling assumptions or data characteristics.

#### Season-level holdout

To test generalization across annual vaccination cycles, we excluded Flu Year 2 Day 0 and Day 7 samples from principal component analysis (PCA) fitting. Subtype-specific centering was recomputed using training data only to avoid leakage of held-out visits into subtype reference profile estimates. The ITL was then refit using the remaining data, and the held-out season was projected into the learned space. Donor-level temporal coherence between Day 0 and Day 7 was quantified using Spearman rank correlation across donors for each principal component (PC) and summarized as the median correlation across PCs 1-20. Temporal coherence in the held-out season was comparable to that obtained from the full model, indicating that the ITL geometry does not depend on inclusion of a specific annual vaccination cycle.

#### Gene program exclusion

As a gene-level sensitivity analysis, we excluded gene sets corresponding to interferon-stimulated genes, immediate early response genes and heat shock genes from the shared gene universe and refit the PCA. This exclusion was not part of the primary analysis, in which such programs were controlled at the cell level through filtering of transient non-homeostatic cells. Multi-dimensional donor-level temporal coherence across PCs 1–20 remained unchanged following exclusion of each gene set, indicating that the ITL is not reducible to any single activation-associated transcriptional program.

#### Aggregation strategy (pseudobulk)

To assess sensitivity to aggregation strategy, we repeated the analysis using a pseudobulk approach in which raw counts were summed within each donor–visit–subtype group prior to normalization. Pseudobulk profiles were normalized by library size, log-transformed, and centered using the same subtype-specific reference profiles as in the primary analysis. PCA was then performed using the same gene universe and analytical pipeline. The resulting ITL exhibited comparable multi-dimensional donor-level temporal coherence across PCs 1–20, indicating that the observed transcriptional structure is not sensitive to the choice of aggregation method.

#### Sequencing depth and complexity

We evaluated the association between ITL coordinates and sequencing complexity metrics computed at the donor-by-visit level, including mean detected genes per cell, mean unique molecular identifiers (UMIs) and mean mitochondrial read fraction. The first principal component exhibited a strong correlation with mean detected genes and UMIs, consistent with a global transcriptional complexity axis. This association persisted within sequencing batch, pool, chip and calendar year strata, arguing against a simple between-batch artifact. Importantly, leading components beyond the first principal component (PCs 2–20) showed minimal correlation with sequencing complexity metrics while retaining strong donor-level temporal coherence.

#### Gene universe threshold sensitivity

To test sensitivity to gene inclusion criteria, we reconstructed the ITL using a relaxed gene prevalence threshold (2% versus 10% in the primary analysis), expanding the shared gene universe from 3,530 to 8,171 genes. Multi-dimensional donor-level temporal coherence across PCs 1–20 remained unchanged, demonstrating that the ITL structure is not sensitive to the specific gene prevalence threshold used for gene universe definition.

#### Cell count sensitivity

To assess the influence of variable group sizes, we downsampled each donor–visit–subtype group to 100 cells prior to aggregation and PCA fitting. Downsampling was performed at the cell level before computation of subtype-resolved mean expression profiles. The resulting ITL retained comparable multi-PC donor-level temporal coherence, indicating that the shared transcriptional structure is not driven by unequal sampling depth across groups.

#### Cross-tissue analyses

Analyses in the cross-tissue immune atlas [17] test whether coordinated donor-level variation identified in the Sound Life cohort is preserved across anatomical contexts and reflected at the level of protein expression. The processed AnnData object provided with the study is used as input. RNA features are restricted to the Sound Life-derived gene universe described above, yielding a partially overlapping gene set (3,425 genes retained in the cross-tissue dataset). Cells are grouped by donor, tissue and native cell type, and expression is aggregated to donor × tissue × cell type mean profiles using the same normalization and aggregation framework applied in the Sound Life cohort, with groups required to meet a minimum cell count threshold (N=100). Deviation profiles are constructed analogously to the Sound Life analysis by centering expression within each cell type and tissue, such that donor-specific deviations are defined relative to the mean expression of that cell type across donors within the same tissue. This removes baseline differences in gene expression associated with tissue context and isolates donor-level variation. For each comparison, analyses are restricted to sets of donors shared between the relevant cell types or tissues, and deviation profiles are computed within these matched donor sets rather than across a global cohort. Only comparisons with at least 10 shared donors are retained.

To evaluate cross-cell type coordination within tissues, gene-wise Spearman correlations are computed between deviation profiles from pairs of cell types within the same tissue and matched donor. To evaluate cross-tissue consistency, correlations are computed between deviation profiles for the same cell type across tissues within the same donor. In both cases, correlations are computed per donor within each matched set and summarized across donors for each comparison unit.

CITE-seq protein measurements are analyzed in parallel using the normalized protein matrix provided with the dataset. Protein expression is aggregated to the same donor × tissue × cell type groups and deviation profiles are constructed using the same within-group centering procedure. Correlation analyses are then performed using identical donor-matched comparisons as for RNA.

Null distributions are generated by permuting donor identities within matched sets, preserving group structure while disrupting within-donor correspondence. These controls test whether observed correlations arise from donor-level alignment rather than shared marginal distributions.

#### Association of the ITL with routine hematologic indices

To assess whether dimensions of the immune transcriptional landscape (ITL) correspond to classical whole-blood physiological measurements, we analyzed their relationship with routine hematologic indices obtained from complete blood count (CBC) testing.

CBC-derived variables were obtained from the Sound Life online dataset. Variables analyzed included hemoglobin concentration, hematocrit, platelet count, total white blood cell count (WBC), absolute neutrophil count, absolute lymphocyte count, absolute monocyte count and corresponding percentage-based measures. Only donor–visit pairs with both ITL coordinates and hematologic measurements available were retained for analysis (*n* = 776).

Associations were evaluated using the reduced ITL manifold representation used in the proteomic correspondence analyses. Donor–visit ITL coordinates were constructed by averaging subtype-specific principal component scores across immune subtypes present in ≥ 95% of donors. Principal component analysis was then applied to the first 150 ITL principal components to obtain a reduced manifold representation. Associations between each ITL manifold coordinate and individual hematologic variables were quantified using Spearman rank correlation.

#### Proteomics manifold construction and ITL-proteomics correspondence analyses

Serum proteomic measurements from the Sound Life cohort were used to assess whether donor-level position within the immune transcriptional landscape corresponds to circulating protein profiles. Proteomic features were derived from a broad panel of circulating proteins measured by Olink, with proteins included based on data availability rather than predefined functional annotation.

Proteomic features were normalized as described in the Sound Life study [19]. Donor–visit protein values were obtained by averaging replicate measurements where applicable. For each holdout analysis, proteins with high number of missing values were excluded using the training data only, and remaining missing values were imputed using training-set medians prior to dimensionality reduction. Principal component analysis was applied to the normalized proteomic matrix to construct a low-dimensional proteomic manifold representation, and the first ten proteomic components were retained for analysis.

To enable cross-modal correspondence analyses, donor–visit ITL coordinates were first constructed by aggregating subtype-level principal component scores across immune subtypes present in ≥ 95% of donors. Because the ITL representation is high dimensional, principal component analysis was applied to the first 150 ITL dimensions to define a reduced ITL manifold. The first ten ITL manifold dimensions were used for correspondence analyses. For each holdout analysis, all preprocessing steps within the proteomic and reduced ITL manifold construction, including feature filtering, imputation and dimensionality reduction, were performed using training data only, and held-out samples were processed using parameters learned from the training data before regression evaluation.

We evaluated the correspondence between proteomic and transcriptional manifolds using ridge regression models mapping proteomic manifold coordinates to ITL manifold coordinates. Model generalization was assessed using two strict holdout strategies: donor holdout (all visits from a subset of donors withheld during training) and temporal holdout (all samples from specific visits withheld). For each analysis, dimensionality reduction and regression models were fit using training data only, and held-out samples were projected into the resulting spaces for evaluation. Model performance was quantified using Spearman correlation between predicted and observed ITL manifold coordinates in the held-out data.

Statistical significance was assessed using permutation testing by randomly permuting ITL targets in the training data, refitting the regression model and evaluating performance on the fixed test set. Empirical *p*-values were computed from the resulting null distribution.

#### Parse perturbation analyses

Analyses of the Parse cytokine perturbation dataset were designed to match the aggregation framework used for the Sound Life cohort while preserving the donor-matched perturbation structure of the experiment. A shared gene universe of 3,438 genes was first defined by intersecting the Parse gene set with the ordered ITL gene list from the Sound Life joint immune landscape. All Parse analyses were restricted to this shared ordered gene set.

Raw UMI count matrices were normalized at the cell level using the same library-size target sum used in the Sound Life analyses (*T* = 4524). For each donor–cell type–cytokine group, mean expression profiles were computed by averaging cell-level log(1 + *x*) normalized expression values across all cells in the group. Only donor–cell type–cytokine groups containing at least 100 cells were retained. This produces a group-by-gene matrix of donor–cell type–perturbation mean expression profiles on the shared ITL gene universe.

To define reference profiles within the perturbation dataset, PBS control groups were identified and averaged within each cell type across all donors. These cell type-specific PBS means were subtracted from all donor–cell type–cytokine group means to produce centered group-level expression profiles. Perturbation response vectors were then defined by donor-matched subtraction of the corresponding PBS-centered group mean within each donor and cell type. Specifically, for each non-PBS donor–cell type–cytokine group, the matched donor–cell type PBS profile was subtracted, yielding a donor-matched perturbation response matrix. Thus, perturbation responses are represented as donor- and cell type-specific deviations from the matched PBS condition, computed after restriction to the shared ITL gene universe.

To assess how strongly perturbation-induced transcriptional changes align with the space of natural immune variation, Parse perturbation response vectors were projected into the fixed ITL basis learned from the Sound Life cohort after restriction to the shared gene universe. The fraction of variance captured by ITL components was then computed from the full perturbation response matrix.

To quantify preservation of donor-level organization under perturbation, within-donor coordination was computed by evaluating pairwise Spearman correlation across cell type-specific perturbation response vectors within each donor in the ITL space (first 150 components) and taking the median across cell type pairs. These within-donor coordination values were then summarized across donors to obtain a single coordination statistic per cytokine. To assess the donor-specific contribution to cross-cell-type coordination, donor labels were independently permuted 1000 times within each cell type for a given cytokine and the coordination analysis was repeated. This permutation disrupts donor matching across cell types while preserving coordination arising from shared responses to each cytokine.

## Statistical analysis

Spearman rank correlation was used throughout to quantify relationships between gene-wise deviation profiles, donor-level transcriptional coordinates and cross-modal features. This choice avoids assumptions of linearity or normality and focuses on preservation of relative ordering across donors.

Where statistical significance was evaluated, null distributions were generated using permutation testing. For deviation alignment analyses, donor identities were permuted within visits while forbidding self-matches. For proteomics correspondence analyses, ITL targets were permuted in the training data prior to model fitting and evaluated on fixed test sets. Empirical *p*-values were computed from the resulting null distributions.

Bootstrap resampling of test samples was used to visualize sampling variability of selected summary statistics. Bootstrap results were used for visualization only and were not used for formal hypothesis testing.

All analyses were performed in Python using NumPy, SciPy, pandas and scikit-learn. Random seeds were fixed where applicable to ensure reproducibility of dimensionality reduction and regression procedures.

## Data Availability

All data used in this study is publicly available. Details of the Sound Life cohort data may be found in Gong et al. [19] and data may be accessed at https://apps.allenimmunology.org/aifi/insights/dynamicsimm-health-age/cohorts/. The Terekhova et al. [16] validation dataset is available via Synapse (syn49637038), the cross-tissue multimodal immune atlas from Wells et al. [17] was accessed via the CZ CellxGene data portal (https://cellxgene.cziscience.com) and the Parse Biosciences cytokine perturbation dataset [18] was obtained from the Parse Biosciences data portal (https://www.parsebiosciences.com/datasets/).

## Acknowledgments

We gratefully acknowledge the Sound Life investigators for generating and making publicly available the longitudinal immune profiling dataset used in this study. The extensive experimental design, longitudinal sampling and multiomic integration underlying this resource made the present analyses possible. We thank Miah Wander, Claire Gustafson and Julia Greissl for providing feedback that improved the manuscript.

## Declaration of generative AI and AI-assisted technologies in the manuscript preparation process

During the preparation of this work the author used ChatGPT 5.4, OpenAI in order to generate code and edit manuscript for clarity. After using this tool, the author reviewed and edited the content as needed and takes full responsibility for the content of the published article.

